# Extreme and rapid bursts of functional adaptations shape bite force in amniotes

**DOI:** 10.1101/333351

**Authors:** Manabu Sakamoto, Marcello Ruta, Chris Venditti

## Abstract

Adaptation is the fundamental driver of functional and biomechanical evolution and can be linked to rates of phenotypic trait evolution. Significant shifts in evolutionary rates are seen as instances of exceptional adaptation. However, whether or not signatures of exceptional adaptations (elevated rates) can be distinguished from general adaptations (background rate) in biomechanical traits remains to be tested in a robust statistical framework. Here, we apply a recently developed phylogenetic statistical approach for detecting exceptional adaptations in bite force, in a large group of terrestrial vertebrates, the amniotes. Our results show that bite force in amniotes evolved through multiple bursts of exceptional changes, whereby whole groups – including Darwin′;s finches, maniraptoran dinosaurs (group of non-avian dinosaurs including birds), anthropoids and hominins (the group of species including modern humans) – experienced significant rate increases compared to the background rate. However, in most parts of the amniote tree of life we find no exceptional rate increases, indicating that coevolution with body size was primarily responsible for the patterns observed in bite force. Our approach represents a template for future studies in functional morphology and biomechanics, where exceptional functional adaptations can be quantified and potentially linked to specific ecological factors underpinning major evolutionary radiations.

## BACKGROUND

Adaptation is the fundamental driver of functional and biomechanical evolution. Measures of biomechanical performance – e.g. bite force – characterize and quantify specific functional performances to fulfil ecological demand – e.g., diet [1, 2]. Functions are therefore typically assumed to be under direct selection – i.e. a change in biomechanical performance indicates selection for changes in function. For instance, taxa with higher bite force are often interpreted to have diets requiring powerful bites driving selection on associated morphological features [3–5]. However, working under the premise that all phenotypic changes are adaptations in one way or another *(general adaptations)* – i.e. phenotypic changes are adaptations regardless of whether or not such changes were under strong direct selection [6, 7] – it would be unjustified to assume that biomechanical traits are proxies for strong selection [8]. Biomechanical traits with strong scaling effects – e.g., bite force – may have simply evolved in tandem with body size [9] and may or may not have been under direct selection. Such cases of general adaptations will invariably still be under selection with changes being proportional to the passage of time – i.e. Brownian motion – but may not be under direct positive selection. Thus, it is important and necessary to distinguish general adaptations from adaptations associated with genuine, strong phenotypic selection, i.e., *exceptional adaptations*.

It is becoming well established that the intensity of natural selection acting on a phenotype is linked to rates of evolution [7, 8], and significant shifts in rates can be interpreted as instances of exceptional adaptations – recently defined as positive phenotypic selection [8]. Positive selection is invoked as an explanation for trait evolution along a branch on a phylogenetic tree if the amount of evolutionary change exceeds the amount of change expected from the passage of time given the background rate of evolution. In the context of biomechanical evolution, evidence for exceptional adaptations in biomechanical traits can be detected as significant rate shifts using this phylogenetic comparative framework. However, in spite of the ever-increasing numbers of comparative biomechanical studies employing a phylogenetic framework [e.g., 10, 11–20], whether or not signatures of exceptional adaptations in biomechanical performance measures can be distinguished from general adaptations has never before been tested in a statistically rigorous and phylogenetic framework.

Here, for the first time, we test the hypothesis that instances of exceptional adaptations have shaped the observed diversity in biomechanical traits focusing on bite force evolution. Our hypothesis specifically relates to detecting exceptional adaptations and not in detecting rate heterogeneity – exceptional adaptation is defined as instances of exceptionally large rate increases (at least twice the background rate, see Methods for details). Bite force relates to species′ niche and feeding ecology, is correlated with several ecological and behavioural traits [21–24], and is widely available from the literature across several fields of study (e.g., biomechanics, ecology, palaeobiology) for a broad sample of the tree of life, making it an ideal biomechanical trait to test our hypothesis using a phylogenetic framework. To this end, we assembled the largest dataset of bite forces collected to date for amniotes, both extinct (including non-avian dinosaurs, saber-toothed cats, and hominins) and extant.

If exceptional adaptations are detected in bite force evolution, then we would expect there to be two types of rate shifts on the phylogenetic tree (Fig. S1). The first type of rate shifts, branch-wise rate shifts (branch shifts), are cases in which significant increases in rates with respect to background rate are detected along individual branches. The second type of rate shift, clade-wise rate shifts (clade shifts), occur across all branches within a clade and represent cases in which rapid divergences in trait values have occurred (Fig. S1).

## METHODS

### Data

Bite force and body mass data were collected primarily through the literature, augmented with novel estimates (Supplementary Material), spanning 434 extant and extinct amniote species (Table S1). We used the Time Tree of Life (TTOL) [26] as the backbone phylogeny, with fossil tips/clades inserted at the appropriate phylogenetic and temporal positions according to current consensus (Supplementary Material) using fossil dates from the Paleobiology Database (accessed 9 Feb 2017).

### Variable-Rate Phylogenetic Regression Models

We fitted a variable-rate regression model [8] using BayesTraits [27] on log10 bite force against log10 body mass *(Single-slope VR model)*. As previous research indicated that scaling of bite force is group-specific [28–30] we tested an additional model in which separate slopes were estimated for five different groups *(5- Group VR model)*. Each taxon was assigned to one of five groups: mammals excluding bats (hereafter “Mammals”), bats (Chiroptera), finches (Passeroidea, the superfamily of songbirds to which Fringillidae, Estrildidae, and Thraupidae [Darwin′s finches] belong); non-passeroid dinosaurs (including other birds, hereafter “Dinosaurs”) and non-dinosaurian reptiles (hereafter “Reptiles”) (Fig. 1; Table S1). We chose these five groups because they act as good descriptors of the distribution of data (Fig. S3) as well as conforming to widely recognized taxonomic groups. Birds and dinosaurs were grouped together, as fossil evidence points to a blurred distinction in their physiology and biology [31–34], while dinosaurs are very different from other reptiles [35]. Intercept differences were not modelled – group-wise offsets will be redundant with rate scalars on branches subtending corresponding phylogenetic groups. In order to account for potential differences owing to bite force acquisition type and biting positions (Supplementary Material), two additional models were fitted with the confounding variables Bite Type and Bite Point individually added to the regression model as covariates (Supplementary Material).

**Figure 1.**
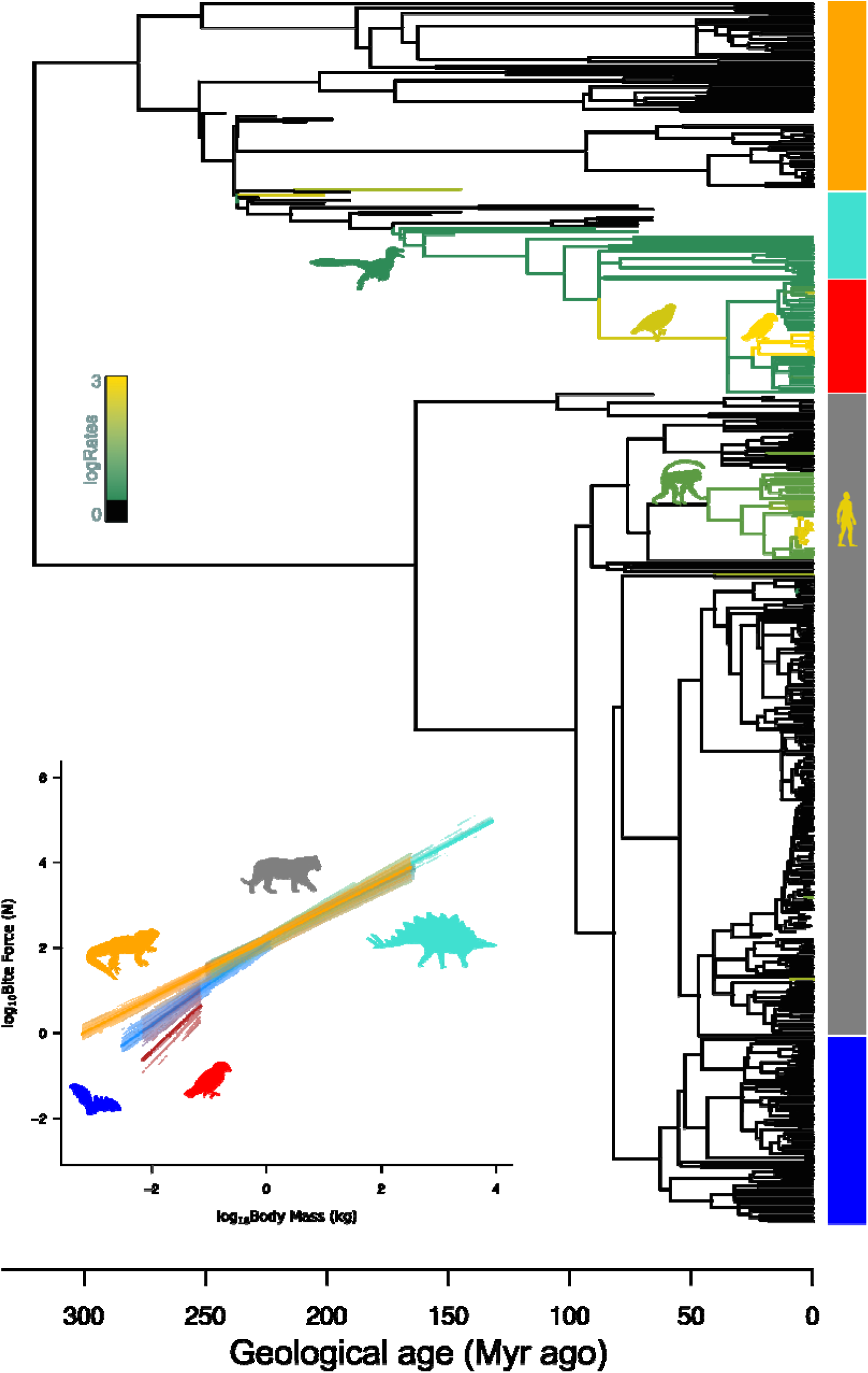
Evolution of bite force and its relationship with body mass. Exceptional rates of bite force evolution are shown as a colour gradient (green to gold) on corresponding branches of the phylogenetic tree used in this study, while branches in which no exceptional adaptations are detected are in black. Silhouettes highlight clades of interest coloured according to corresponding rates: dromaeosaur, Maniraptora; *Platyspiza*, branch subtending Passeroidea; *Geospiza fuliginosa*, Thraupidae; Papionin monkey, Anthropoidea; and *Homo sapiens*, hominin lineage. Inset, the fitted regression lines from a 5-Group variable-rate regression model accounting for bite point (5-Group+BitePoint VR model) across all MCMC runs are shown in colours corresponding to the five groups of interest: bats, blue; mammals excluding bats, grey; finches, red; dinosaurs excluding finches, turquoise; and reptiles excluding dinosaurs, orange. Significant differences in slopes do not exist between bats and finches as well as between mammals, reptiles and dinosaurs, but significant differences exist between the two sets of groups – i.e., bats/finches and mammals/reptiles/dinosaurs (Table S3). Similarly, slopes in bats and finches are significantly different from the theoretical slope of 0.67 but those in mammals, reptiles and dinosaurs are not (Table S4).

We tested for instances of exceptional bursts of evolutionary change in bite force based on rate shifts along branches on the phylogeny [8]. The VR regression model as implemented in BayesTraits works to modify the branch lengths to “detect heterogeneity in rates of phylogenetically structured residual errors” [8]. That is, once the appropriate level of variance in the response variable – e.g., bite force – is explained by some predictor variable(s) – e.g., body mass – outlying deviations from the regression line will be explained as rate shifts (Fig. S1). Under Brownian motion, bite force – after accounting for body mass and other confounding variables – evolves at a rate proportional to time (and an estimated background variance) across the phylogeny, and for any evolutionary change along a given branch that is greater/less than the expected amount of change for the duration of time to occur (given body mass), that branch must be stretched/compressed in length in proportion to the observed amount of phenotypic change – corresponding to a rate increase/decrease. The magnitude of branch stretching/compressing is the rate scalar (*r*). It follows that, phenotypic changes owing to adaptations (potentially as a response to strong selective pressure) would be proportional to *r*. Thus, we define exceptional change following the criteria of [8]: 1) certainty of rate shifts, the branch in question must be scaled in >95% of the posterior sample of scaled trees; and 2) magnitude of rate shifts, the r in question must be greater than two. Rate heterogeneity that do not fulfil these two criteria were not considered as instances of exceptional adaptations. We determined whether rate-shifts constituted exceptional adaptations if they satisfied the criteria set out above in all of three independent replicate Markov Chain Monte Carlo (MCMC) chains.

In order to determine if rate-heterogeneity was statistically significant, we fitted an *equal-rate (ER) model* (or Brownian motion) as a simpler alternative to each of our VR models. Model selection was performed using the Bayes Factor (BF): BF is defined as twice the difference in log marginal likelihood *(m)* between the complex model (model_1_) and the simple model (model_0_) – i.e., BF = 2 × *(m*_1_ – *m*_0_). For instance, we computed BF using m from our 5-Group VR model and the simple alternative 5-Group ER model, and selected the VR model over the ER model when BF value was greater than 2 [36].

We ran our MCMC chains for 10^9^ iterations, with a burn-in period of 10^8^ iterations, sampling every 10^5^ iterations, resulting in a posterior sample of 900 modified VR trees and model estimates, for each regression model. We used stepping stone sampling to compute marginal likelihoods from which BF were calculated. Post-processing of the BayesTraits outputs were conducted using an online post-processor (available at www.evolution.reading.ac.uk/VarRatesWebPP), as well as in R [37].

## RESULTS

### Variable-Rate Regression Model

We found strong support for the VR model compared to the ER model (BF_VR-ER_ = 474) for the single-slope regression model. Bite force scales nearly isometrically with body mass, with a slope of 0.674 (*p*MCMC_0_ < 0.001, R^2^ _mean_ = 0.79; Table S2), which is not significantly different from a theoretical isometric slope of 0.67 (*p*MCMC_0.67_ = 0.4) [38].

There is statistical support for favouring the 5-Group model over the single-slope model (Fig. 1)); significant differences exist among the slopes of different groups (Table S2; Table S3). Finches and bats are not different from each other, but are distinct from mammals, reptiles and dinosaurs (Fig. 1; Table S3); in turn, these three groups are not different from each other (Fig. 1; Table S3). Finches and bats have slopes that deviate from 0.67 (Table S4), while the other three groups have slopes that are not significantly different from 0.67 (Table S4). Critically, despite allowing for the variation in slopes among taxonomic groups, our 5-Group VR model (R^2^ _mean_ = 0.809; Table S2) still outperforms a 5-Group ER model (BF_VR-ER_ = 429; Table S2).

Model selection showed that bite type is not significant (*p*MCMC > 0.05 in both Single-Slope+BiteType and 5-Group+BiteType models; Table S2) while bite point is (*p*MCMC < 0.05 in both Single-Slope+BitePoint and 5-Group+BitePoint models; Table S2). Given a similar body size, bite force is comparable in magnitude between *in vivo* measurements and indirect estimates, but it differs in magnitude between posterior and anterior bites (posterior positions have higher forces, as expected). There is no slope difference between bite type categories or between bite point categories (Figs S6, S7). We used the 5-Group+BitePoint model as our final model for detecting exceptional adaptations in bite force – this enables us to compare evolutionary rates after accounting for effects owing to body size and bite point.

### Rate Shifts and Exceptional Adaptations

We found substantial amount of rate heterogeneity (elevated rates in >50% of the posterior sample) in the amniote tree of life, along 439 branches out of 866 branches in the phylogeny (51% of branches; Fig. S4). Instances of exceptional adaptations are found in a far fewer number of branches: in 182 branches (21% of branches) (Table S5; Figs 1, 2a, S5; Movie S1). Our results show that bite force evolved through multiple bursts of exceptional adaptations, whereby whole groups experienced rate increases of bite force evolution compared to the background rate across the entire amniote tree. We find such clade shifts in Darwin′s finches (median r > 55), the hominin lineage excluding *Australopithecus anamensis* (*r* > 35), Anthropoidea (*r* >6), and maniraptoran theropod dinosaurs (the clade including birds and their closest relatives, here *Erlikosaurus and Dromaeosaurus; median r* > 3).

**Figure 2.**
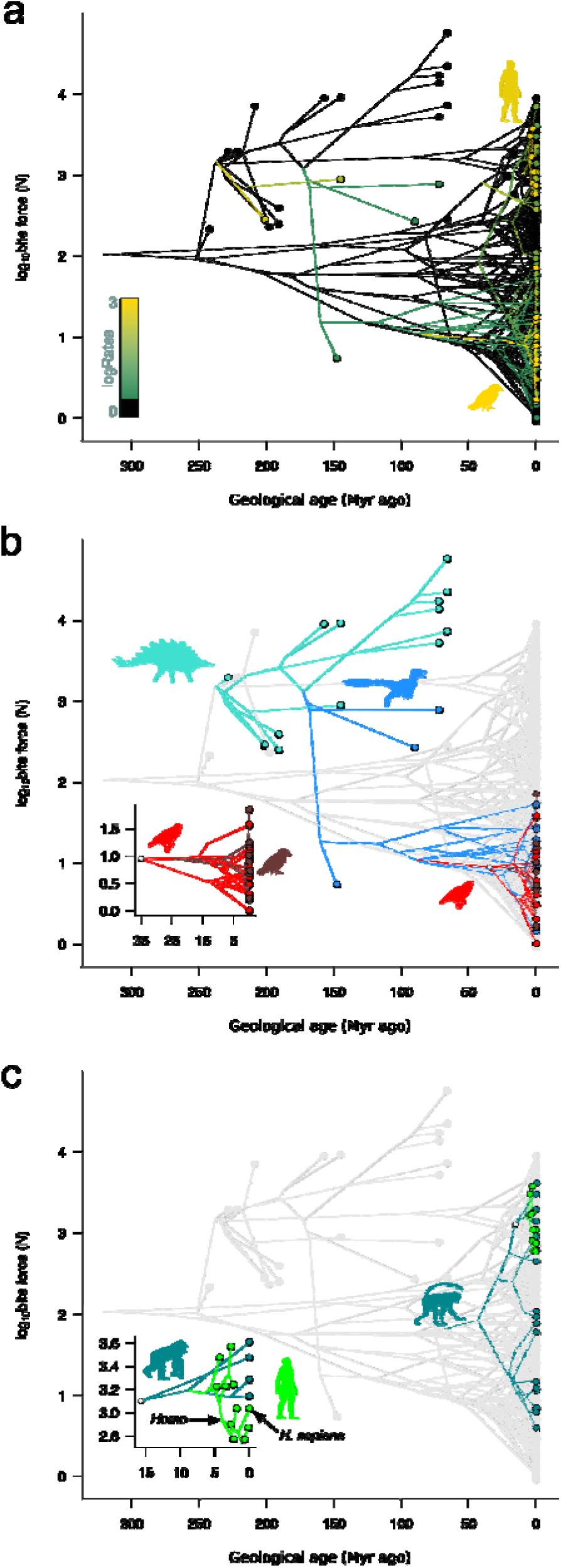
Ancestral reconstruction of bite force across phylogeny and through time. Evolution of bite force with respect to body size while accounting for variable rates show branches and clades with higher amount of change in bite force than expected given ancestral body sizes and phylogenetic positions. Exceptional rates along the branches of the whole tree are shown as a colour gradient (green to gold), with the two clades exhibiting the highest rates (Darwin’s finches and hominins) indicated with silhouettes coloured according to corresponding rates (a). Clades in which exceptional rates were detected are highlighted: b, Dinosauria (turquoise) with maniraptorans (skyblue), finches (red) and Darwin’s finches (brown); and c, Anthropoidea (teal) and hominins (bright green). Insets show subclades of interest (node denoted by a white circle in the whole tree): Darwin’s finches amongst the Passeroidea (b) and hominins amongst Hominidae – i.e., great apes (c).

Two aspects of the distribution of rate shifts are noteworthy. First, clade-wise rate shifts show a nested pattern (Fig. 1). For example, Darwin′;s finches exhibit an additional level of rate increase above that of the maniraptoran rate increase (Figs 1, 2a, 2b, S5; Movies S1-S2). The same pattern characterizes hominins (*excluding A. anamensis*) within anthropoid primates (Figs 1, 2a, 2c, S5; Movies S1-S2). In particular, Homo species exhibit reductions in bite forces (Fig. 2c), which is in marked contrast to the apparent increase in hominin body size through time (Figs S8) [39]. Thus, humans drastically reduced bite force through time at a rate faster than their anthropoid ancestors and relatives.

The second key aspect of the rate shift distribution is that branch-wise rate increases occur in conjunction with clade-wide shifts. We identified an exceptional increase in the rate of bite force evolution along the branch leading to Passeroidea (median *r* > 30), followed by a reversal to the ancestral maniraptoran rate. We also recovered a scattering of branch-specific shifts on terminal branches (*Proteles cristatus, Panthera onca, Sus scrofa, Stegosaurus, and Plateosaurus*), marking sudden changes in the biomechanical performance of some species from their close relatives (Figs 1, 2a, S5; Movie S1).

## DISCUSSION

### Exceptional Adaptations in Bite Force

Working under the premise that the rate of phenotypic trait evolution is proportional to the strength of selection [7, 8], we provide the first evidence for exceptional adaptations shaping the diversity of bite force in both extinct and extant amniotes, using a statistically robust evolutionary framework [8]. We find such instances of exceptional adaptations in four clades of amniotes and a handful of independent branches. Conversely, in most lineages of amniotes studied here (79% of branches in the phylogeny), bite force does not undergo exceptional adaptations (even in lineages with elevated rates; Supplementary Material; Fig. S4), indicating that co-evolution with body size is, for the most part, the main factor responsible for bite force variation. A large predator can generate enough bite force to kill its prey just by being large. As an example, *Tyrannosaurus rex* was most likely capable of “pulverizing” bones [40] simply owing to its colossal size (∼ 5-10 tonnes [41–43]). We did not detect instances of exceptional adaptations in bite force in this taxon, and therefore there is no evidence of strong selection for a feeding ecology that requires disproportionately high bite force – e.g. “extreme osteophagy (bone consumption)” [40]. Similarly, we do not detect signatures of exceptional adaptations in classically recognised power-biters such as osteophagous hyenids [44], short-faced hyper-carnivorous felids [45, 46], and small-brained carnivorous marsupials [3, 5, 47] indicating that bite force in these clades were not subjected to strong selection.

Interestingly, contrary to our prior expectations, we do not detect exceptional adaptations in sabre-toothed cats (Machairodontinae). Although rates are on average higher in Felidae as a whole (including both sabre- and conical-toothed cats along with the basal cats *Proailurus* and *Hyperailurictis*) compared to the background rate in the majority of the posterior sample (i.e., >50%; Supplementary Material; Fig. S4), they do not fulfil the criteria for exceptional adaptations. Further, there is no difference between rates in machairodontine lineages and other felid lineages. The time elapsed in the lineage leading to sabre-toothed cats since their divergence with conical-toothed cats sufficiently explains the reduction in bite force in sabre-toothed cats.

It is possible that for the majority of taxa in our bite force sample, individuals preferentially seek out and consume food items that can be processed within the naturally generated range of bite forces, and rarely actively seek food items that require maximum biting capacity. This equates to a behavioural adaptation, in which species evolve bite force through correlated evolution with body mass, and preferentially consume food items that fit within their natural range of bite force. If this is true, then selection for improved biting performance with respect to higher bite force may not frequently occur. Similarly, drastic reduction in bite force – as a trade-off between force and velocity if jaw closing velocity was under strong selection – would constitute an exceptional adaptation significantly below the expected range of bite force given the universal scaling relationship. However, it is potentially more likely for reductions in bite force to occur over exceptional gains in bite force since muscles are expensive organs to maintain and more so to enlarge. Indeed, we observe more instances of drastic reductions in bite force associated with exceptional adaptations than we do exceptional increases in bite force (Figs 2, S5).

Despite the overall uniform evolution of bite force relative to body mass, we find evidence for exceptional adaptations playing a major role in the evolution of bite force (just not as a tree-wide pattern across all major clades). Since the majority of these exceptional adaptations occur as clade-wide rate shifts, it is possible that they are linked to some biological, ecological or environmental features unique to those clades and shared amongst constituent members. For instance: the acquisition of a “key” innovation, which allows such clades to rapidly expand and exploit functional niches; a shift into a new environment, habitat or lifestyle that is associated with new opportunities and resources; or an extrinsic environmental event (such as mass extinction events) that results in an abundance of ecological niches available for exploitation. Determining such factors ultimately responsible for exceptional adaptations is theoretically possible. Namely, the VR regression framework allows for the inclusion of extrinsic factors such as dietary preference, feeding strategy, sexual display/conflict, etc., as additional covariates. A covariate can be identified as the extrinsic driver of bite force evolution if its inclusion can explain much of the variation in bite force, thereby reducing or eliminating rate shifts. At present, ecological data associated with biting performance are only available for a handful of species, but we hope that future work will considerably augment information on ecological covariates.

### Macroevolutionary Patterns of Exceptional Adaptations

Overall, our results highlight a combination of clade-wise and branch-wise rate shifts occurring across the amniote phylogeny. Clade-wise rate shifts are characterized by an elevated rate that is homogenous across all branches within a given clade and are associated with an increase in trait variation in the constituent taxa given the variance in the other taxa in the data [8]. Lineages in such clades continually evolve traits at a faster rate through time compared to other parts of the tree (Fig. 1). This contrasts with a classic description of an adaptive radiation [48], which is characterized by a rapid initial burst of trait evolution followed by a rate slowdown associated with niche saturation [49]; clade-wise rate shifts see continual changes in functional niche occupation. One implication of such patterns in bite force evolution is that evolutionary lineages (sequence of branches leading to terminal taxa) do not stay in the same regions of function-space (pertaining to biting functional variation), but rather, continue to expand out to unoccupied/unexplored regions of function-space. Evolutionary lineages will be moving through various functional niches as their bite force values change through time. An alternative interpretation is that function-space itself changes through time. i.e. functional/ecological niches are dynamic rather than fixed entities, a constantly moving target [50]. Yet another interpretation is that function-space saturates but convergences occur frequently and rapidly – that is, lineages move in and out of occupied/explored regions of function-space. In this context, our results would support the notion that functional adaptations are relatively labile over evolutionary history and remain responsive to changing environmental conditions and ecological demands.

Branch-wise rate shifts (rate shifts associated with single branches only) occurring on branches subtending whole clades (Fig. 1), such as that observed at the base of Passeroidea in our dataset, can be interpreted as a mean-shift in bite force after accounting for body mass [8]. In our case, this means that there was a rapid shift in the mean bite force value of Passeroidea from the ancestral maniraptoran mean. The total sum of evolutionary changes accumulating along the branch leading to Passeroidea exceeds that expected from the temporal duration of that branch. This is irrespective of any un-sampled taxa along the lineage – e.g., other perching birds (Passeriformes, e.g. corvids, shrikes) for which bite force data are not available in the literature as far as we are aware.

Similarly, the two large-bodied herbivorous taxa, *Stegosaurus and Plateosaurus*, have evolved bite force at excessively high rates (∼ 11 and ∼ 35 times background rate, respectively – *Stegosaurus* and *Plateosaurus* have extremely small heads, and thus low bite forces, for their body sizes), but these could potentially represent evolutionary patterns within thyreophoran and sauropodomorph dinosaurs respectively, and not specifically associated with these two species. Nonetheless, major changes in bite force did occur along these lineages so the interpretation remains the same: the amount of change in trait values given the duration of time elapsed is exceptionally high compared to the background rate.

### Evolution of Bite Force in Dinosaurs and Birds

Maniraptoran theropods are perhaps the most diverse amongst dinosaurs in terms of functional and morphological specializations associated with feeding. Forms like the parrot-like oviraptorosaurs, large herbivorous therizinosaurs, hyper-carnivorous dromaeosaurs with recurved teeth (e.g. *Velociraptor*), and toothed and toothless avialans are just some typical examples of maniraptoran morpho-functional diversity. High evolutionary rates in maniraptoran bite force indicate that their morphological and presumed ecological diversity are linked with selection on biting performance. Maniraptoran fossils are predominantly known from Cretaceous rocks but are inferred to have originated by the Middle Jurassic (∼ 168 Myr ago; Fig. 1), with derived members including the avialan *Archaeopteryx* appearing relatively quickly, by the Late Jurassic (∼ 150 Myr ago; Fig. 1). This implies that Maniraptora underwent a rapid diversification (both in species diversity but also in bite force variance) early in their evolutionary history, but that they retained high evolutionary rates in bite force throughout the clade’;s history.

The observation that bite force underwent exceptional adaptations in maniraptoran theropods but not uniquely in birds – rates in birds are not distinguishable from those in other non-avian dinosaurs – is consistent with recent findings that the evolution of birds and their immediate close relatives – i.e. paravians – are similar to one another [31], and that many of the features traditionally associated with birds were present in paravians and more broadly in maniraptorans. Here we have demonstrated that this is also the case with bite force evolution (given the available data); the rate of bite force evolution did not change from non-avian maniraptorans to birds. On the other hand, this means that heritable rates of bite force evolution in maniraptoran ancestors possibly contributed to some extent on the subsequent ecological success of birds – the ability to rapidly change bite force in response to changing environmental and ecological pressures would surely have been beneficial for early Cenozoic birds in the post-extinction world.

Our identification of an extreme clade-wide rate shift in the Darwin’s finches, which is among the highest in the tree (>55 times the background rate; Figs 1, 2, S5; Movies S1-S2; Table S5), is noteworthy for both historical and biological reasons. Darwin’s finches are the classic textbook case of ‘adaptive radiation’, with eco-morphological diversification occurring in a short time interval after the initial colonization of the Galapagos Islands by finches [51–53]. Their diversification in feeding ecology is particularly relevant to the rapid evolution of bite force, as Darwin’;s finches are well documented to have strong dietary preferences on food types of varying toughness [54] or differences in food manipulations [55]. Within the context of the evolutionary history of amniotes as sampled here (∼ 350 Myr), the radiation of Darwin’s finches is comparatively recent with some divergences occurring in a geologically instantaneous manner (Fig. 1). Compared to their recent divergence times, bite force variance in Darwin’s finches is exceptionally high spanning almost two orders of magnitude (Fig. 2) accounting for their extraordinarily high evolutionary rates.

### Evolution of Bite Force in Humans

The exceptionally high rates of bite force evolution in the hominins excluding *A. anamensis* (Figs 1, 2) – more than 35 times the background rate and ∼ 6 times those for the branches within other anthropoid primates – highlights an important, recent and rapid evolution in our own lineage. In particular, the decrease in bite force in *Homo species* (Fig 2) is contrary to the increase in hominin body size through time (Fig S8) – such a discrepancy between bite force and body mass is indicative of strong directional selection, and coincides with previously documented evolutionary shifts in relative molar sizes, attributed to the reduction in feeding time associated with the introduction of food processing such as cooking [8, 56].

Additionally, the reduction in bite force in the hominin lineage may have occurred as a consequence of an evolutionary trade-off with increasing brain size in this group [57–59] (see Supplementary Material; Figs S9-S10). As brain size increases relative to skull size, the temporal fossa (defined as the opening between the braincase and the zygomatic arch) is reduced in dimension, thereby decreasing the amount of space available to house the temporal muscles [58, 59], which are critical for achieving hard biting in most animals. This reduction in temporal muscle can be seen in the changing predominance of the sagittal crest (ridge of bone running along the midline of the skull) where the temporal muscles attach through hominin evolution (Fig. S10). Strikingly, molecular evidence supports the hypothesis that a drastic reduction in the temporal muscle occurred along the lineage to *H*. *sapiens* after its divergence from chimpanzees owing to a frameshifting mutation, causing inactivation of the predominant myosin heavy chain expression in masticatory muscles [58]. The mutation has been inferred to have coincided approximately with the enlargement of the brain, presumably concurrent with the origin of the genus *Homo* [58] – though the timing has been contested [60, 61]. Furthermore, *H. sapiens* relies less on the temporal muscles and more on the masseter muscles to generate bite force [62]. Thus, such a trade-off between temporal muscle size and brain size is a reasonable explanation for the evolutionary reduction in bite force in the hominin lineage through time, with the advent of cooking further accelerating the loss of reliance on high bite force for food processing. Indeed, an auxiliary phylogenetic regression modelling [63] of bite force on brain size [endocranial volume; 64, 65, 66] accounting for body mass on hominids (*Pongo*, *Gorilla*, *Pan*, and hominins), shows that bite force scales negatively with brain size (Supplementary Material; Fig. S9). The reduction in bite force is statistically associated with an increase in brain size.

## CONCLUSIONS

Taken together, our results reveal that the evolution of bite force in amniotes occurred as bursts of accelerated changes across multiple clades, and as the product of repeated and nested pulses of progressively higher rates of change, representing instances of exceptional functional adaptations. Using a phylogenetic evolutionary framework, on a dataset representing the largest taxonomic sample to date, enables us to statistically detect instances of adaptations in biomechanical metrics, not only along specific branches, but also through time, paving the way to better understand how specific ecological niches (feeding ecologies) are occupied. In order to determine whether species’; bite force underwent instances of exceptional adaptations, it is necessary to demonstrate an exceptionally high rate of evolution associated with that species in a phylogenetic context after accounting for size and the expected evolutionary change associated with divergence time.

## SUPPLEMENTARY MATERIAL

Data are available as Electronic Supplementary Material and on the Dryad Digital Repository (doi:10.5061/dryad.q12c06f).

## ACKNOWLEDGMENTS

We thank Joanna Baker, Ciara O’;Donovan, Andrew Meade, Jorge Avaria Llautureo, and Henry Ferguson-Gow for invaluable discussion, which helped improve the research and manuscript. MS thanks the curators and collection managers at the various institutions visited as part of the data collection process, in particular Rhian Rowson (Bristol City Museum and Arts Gallery), Paolo Viscardi (National Museum of Ireland – Natural History; formerly Horniman Museum), Mark Carnall (Oxford University Museum; formerly Grant Museum, UCL), Lars Werdelin (Swedish Museum of Natural History), Andrew Kitchener (National Museums Scotland – Natural Sciences), Judy Galkin (American Museum of Natural History), Roberto Portela Miguez (Natural History Museum, London), and Milly Farrell (Oxford Brookes University; formerly Hunterian Museum, Royal College of Surgeons). We also thank Larry Witmer (Ohio University) for permission to use his gallery of various theropod skull images. Finally, we thank the reviewers for their constructive comments on this manuscript. Silhouettes in Figs 1 and 2 are attributed to Andrew Farke (*Stegosaurus*, CC-BY 3.0), Michael Keesey (*Homo sapiens*, CC-BY 3.0; *Gorilla*, CC0 1.0), Steven Traver (*Sphenodon punctatus*, CC0 1.0), Sarah Werning (*Panthera tigris*, CC-BY 3.0) and Yan Wong (Chiropteran, CC0 1.0). Anthropoid is uncredited (CC0 1.0). Silhouettes of finches were drawn using photographs by Vince Smith (Geospiza fuliginosa, CC-BY 2.0) and Brian Gratwicke (*Platyspiza crassirostris*, CC-BY 2.0). Silhouette of dromaeosaur is author’s own work (MS).

